# Physics-Driven Zero-Shot Reconstruction of Isotropic 3D Fluorescence Microscopy under Undersampled Acquisition

**DOI:** 10.64898/2026.06.13.732100

**Authors:** Ruijie Cao, Tong Jin, Fengyuan Xin, Yiwei Hou, Yunzhe Fu, Boya Jin, Lijun Li, Shu Gao, Han Wang, Yaning Li, Dilizhatai Saimi, Wei Ren, Wenyi Wang, Guangwei Xin, Kexin Yuan, Zhixing Chen, Xuantao Su, Donghyun Kim, Meiqi Li, Peng Xi

## Abstract

Three-dimensional (3D) imaging represents the development of next generation of fluorescence microscopy. However, routine axial down-sampling makes isotropic resolution unrealistic. Here, we propose DeepUI, a physical zero-shot framework designed to achieve isotropic 3D fluorescence images from a low axial sampling rate. DeepUI fully leverages the intrinsic characteristics of 3D images through physics-guided degradation, which incorporates spatial-frequency joint learning to generate a scaled optical transfer function, combined with noise degradation and an up-sampling branch. Typically requiring just 5 minutes for training and 0.5 minutes for high-throughput and fast prediction, we demonstrate the superior performance of DeepUI to get isotropic results, and the exclusivity to axial down-sampling conditions, even in more challenging conditions, including defocused background, noise, and resolution blur.

## Introduction

Three-dimensional fluorescence microscopy is a central tool for resolving the structural organization and dynamic processes of biological systems^1-4^. In principle, volumetric information is obtained through axial sampling; however, the axial resolution remains substantially lower than the lateral resolution due to physical constraints, leading to the long-standing anisotropy resolution^5-10^.

Beyond this well-recognized anisotropy resolution, a more fundamental challenge arises in practice: the mismatch between acquisition strategies and computational reconstruction^11,12^. While dense axial sampling is required to approach diffraction-limited resolution, it imposes severe compromise on imaging speed, phototoxicity, and data volume. Consequently, axial under-sampling is widely adopted in real experiments, particularly in live-cell, in vivo, and high-throughput imaging, resulting in substantial deviations from the Nyquist sampling condition and significant loss of axial information.

Existing solutions attempt to mitigate anisotropy either through advanced optical designs or computational reconstruction. Optical approaches, such as dual-objective or interference-based systems^13-18^, can partially improve axial resolution but require complex instrumentation and still rely on dense sampling. On the computational side, methods including classical deconvolution such as ^19-22^, supervised learning^23, 24^, and self-supervised restoration^25,26, 27^ have been proposed to enhance axial resolution.

However, we find that these computational approaches share an implicit assumption: the input data are close to fully sampled or only slightly under-sampled. Under realistic under-sampling conditions, this assumption is violated. As a result, existing methods systematically fail, producing structural hallucinations, periodic artifacts, or degraded reconstructions that are insufficient for quantitative biological analysis. This reveals a fundamental mismatch between current algorithmic designs and the data regimes encountered in routine microscopy.

To address this challenge, we reformulate isotropic reconstruction explicitly under under-sampled acquisition. We introduce DeepUI, a physics-driven zero-shot framework that reconstructs isotropic 3D information directly from under-sampled inputs. By embedding the full physical image formation process— including optical transfer function scaling, noise statistics, and axial sampling—into a self-supervised optimization framework, DeepUI eliminates the need for isotropic ground truth or densely sampled training data. This formulation enables stable, high-fidelity reconstruction from strongly under-sampled measurements and reframes under-sampling from a practical limitation into a computationally invertible acquisition strategy, establishing a new regime for 3D fluorescence microscopy.

## Results

### Principles and results of DeepUI

A fundamental principle is that biological specimens are inherently isotropic—they possess no intrinsic orientation preference relative to the observer. The anisotropy observed between lateral and axial dimensions stems solely from the imaging system, specifically the anisotropic PSF, where axial information is captured within a narrower angular range. To achieve isotropic *xyz* imaging, the core idea of DeepUI is to first emulate this axial degradation: we artificially degrade isotropic *xoy* images to simulate anisotropic *xoz* (or *yoz*) views. These degraded images then serve as input, while the original isotropic *xy* images act as the ground truth. A deep learning model is trained to learn this anisotropic transformation, which is subsequently applied along the *z*-axis to generate fully isotropic 3D reconstructions. Traditional methods usually use the generative adversarial network (GAN)^26,27^ or simulated ways^25,28^ to degrade the *xoy* image. However, we demonstrate the concerns of those methods in terms of application scenarios and restoration artifacts. As a result, we integrated the physical degradation model to complete the degradation. We proposed a scaled optical transfer function (*sOTF*) to determine the resolution relationship between *xoy* and *xoz* images. Assuming the frequency domain of *xoz* images can be expressed as *freq*_*xoz*_ = *freq*_*xoy*_ · *sOTF* , and the simulated *sOTF* can be expressed as:

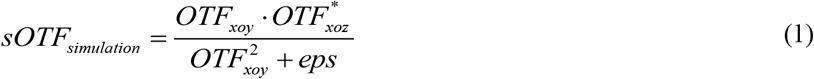

Where *OTF*_*xoy*_ and *OTF*_*xo*z_ means the optical transfer function in *xoy* and *xoz* planes. * means the transposition symbol, and *eps* is the extremely small quantity to avoid a zero denominator. The *sOTF*_*simulation*_ is calculated based on the physical parameters and existing 3D OTF model^29,30^, then 3D anisotropic images are sectioned into small blocks of *xoy* and *xoz* image batches for the degradation. Considering the simulated OTF usually fail to correspond well with experimental OTF, causing insufficient resolution improvement and reconstruction artifacts. So, we adopt a dual-constraint *sOTF* optimization method in Fig. 1(a). To increase the information density, every batch of *xoy* and *xoz* images is summed up to form sum *xoy* images 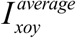 and *xoz* images 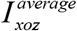 with rich sample information. Then, the frequency domain is obtained using Fourier transform. The circular *xoy* frequency 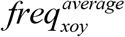 and ellipse-like *xoz* frequency 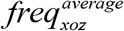 are obtained. At last, the *sOTF* is applied to multiply the *xoy* frequency, and the multiplied results 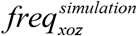 need to approach the *xoz* frequency 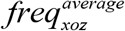. And the *sOTF* is also constrained by the spatial difference between the simulated and calculated *sOTF*. The dual constraint of the spatial and frequency domains can ensure the correctness of calculating the *sOTF*.

**Figure 1.**
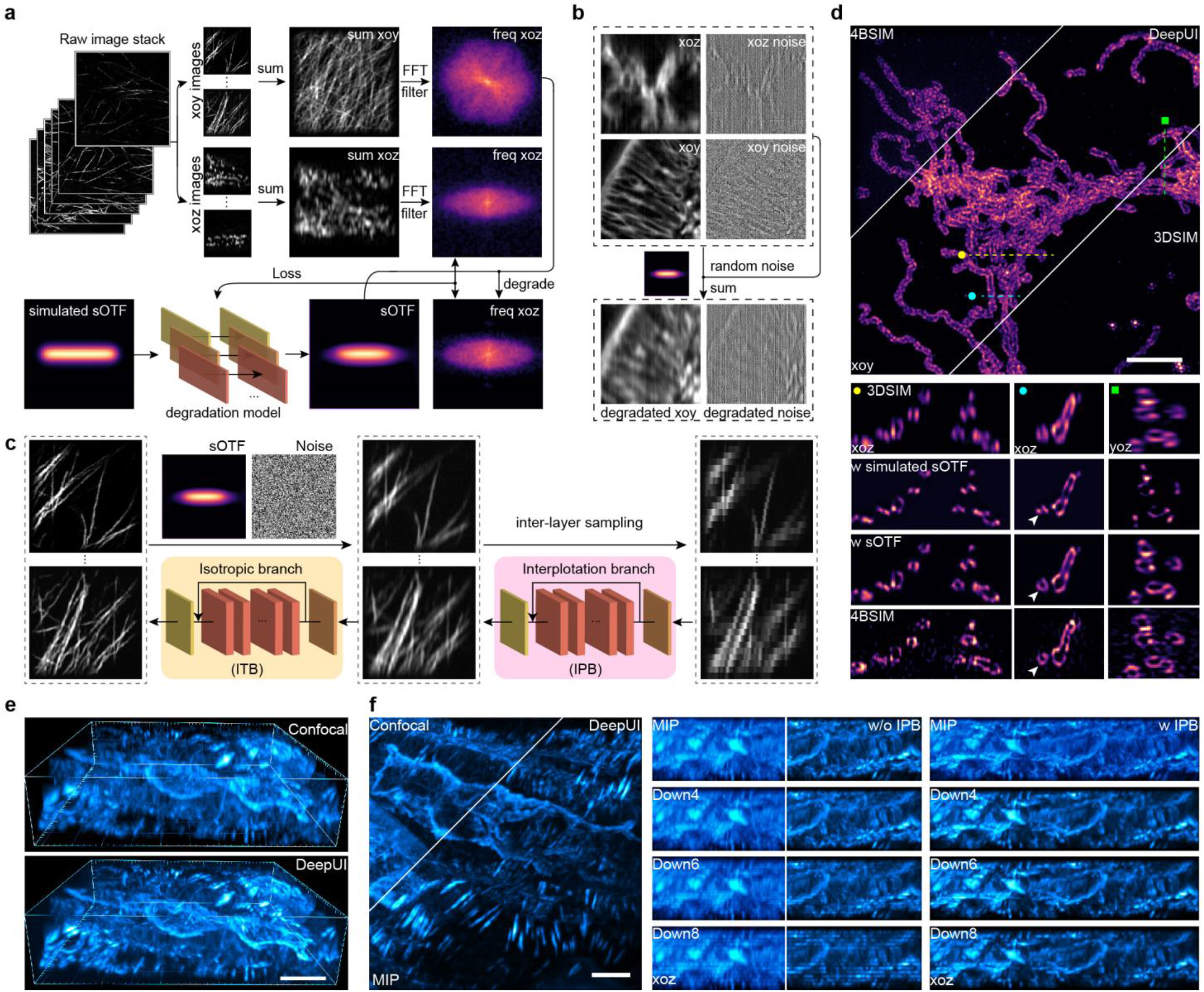
Principles of the degradation model and restoration steps of DeepUI with typical performance. The pre-trained degradation of (**a**) *sOTF* and (**b**) noise. After this process, *sOTF* and noise distribution can be obtained. (**c**) Training process of DeepUI. Paired high/low resolution images (ROI_1_∼ROI_n_) can be obtained, and DeepUI can be trained. (**d**) Isotropic performance of mitochondria imaged by 3DSIM, using 4BSIM as ground-truth. And the comparison of DeepUI using simulated OTF and calculated *sOTF* is presented. (**e**) The 3D view of the mouse kidney actin with Confocal and DeepUI results. (**f**) The performance comparison of DeepUI at different axial scanning sizes of 2, 4, 6, and 8-fold than the lateral pixel size. Scale bar: 4 μm.

Another issue for the degradation is the noise in the *xoy* and *xoz* planes because various noise commonly exists in the fluorescence 3D images. Considering the pixel size in the lateral plane is usually smaller than the axial sampling size, the noise in *xoz* and *xoy* planes will exhibit prominent differences because of the up-sampling operation. So, we firstly use a Gaussian blur to extract the noise-free part of the image, and extract the high-frequency noise for random matching. To increase the stability of the network, we also added a little random noise in the noise part. So, the simulated *xoz* images based on paired *xoy* images can be expressed as:

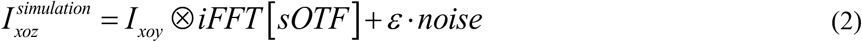

Where *iFFT* [·] means the inverse Fourier transform, *ε* is the parameter to control the noise degree. It can be seen in Fig. 1(b), the simulated *xoz* images have vertical noise rather than random noise in *xoy* images. This can better approach the real *xoz* images.

After the degradation, the paired images can be obtained. The isotropic *xoy* images are degraded with *sOTF* and noise following a strict physical process in Fig. 1(c), and the simulated anisotropic *xoz* images can be obtained. To resolve the situations where axial planes are down-sampled severely, we inter-layer sample degraded *xoy* images along one direction for further degradation. Then, a two-step network is used to restore the isotropic result of the degraded image, where the interpolation branch (IPB) is used to interpolate along the axial direction and the isotropic branch (ISB) is used to restore the isotropic results. And in situations where down-sampling is not severe, ISB without IPB is enough to be used, which can reduce the training and prediction time. To this step, the training process is finished, and the two-step network follows the residual channel attention network (RCAN) structure^31,32^. We section every *xoz* and *yoz* images to conduct the interpolation and isotropic restoration. And the two restored stacks are averaged to form the final result. This prediction process traverses the 3D network into the 2D process and will greatly accelerate the prediction process.

The results of DeepUI to restore the isotropic outer membrane of mitochondria are shown in Fig. 1(d). The comparison of three-dimensional structured illumination microscopy (3DSIM), 3DSIM processed by DeepUI, and 4BSIM (as GT) is shown. And DeepUI with simulated *sOTF* and with calculated *sOTF* through dual-supervised are shown in the *xoz* and *yoz* plane. It can be seen that DeepUI with simulated *sOTF* is different from 4BSIM with extensive sharpening and loss of weak information. But DeepUI with calculated *sOTF* shows a hollow circular structure of the mitochondria outer membrane, showing great similarity to 4BSIM results.

We further use the actin filament with confocal microscopy to showcase the result of DeepUI in Fig. 1(e). In this 3D view, the blurred details can be greatly restored with a greatly improved axial resolution with the help of DeepUI. In 3D images, the layer-by-layer scanning is time-consuming and may cause photon damage. In DeepUI, the interpolation branch (IPB) will solve this limitation to a great extent through deep learning. When the scanning size ranges from 4 to 8-fold compared with the lateral pixel size, namely axial scanning distance of 440 to 880 nm, the DeepUI results without IPB become gradually different from the full-sampled results (as GT) with the loss of axial information and line-shaped artifacts. On the contrary, DeepUI with IPB can greatly reduce the non-continuity of the *xoz* image even with severe down-sampling size in Fig. 1(f, g). So, it’s reasonable that DeepUI can reduce the sampling size of axial planes to a great extent, bringing the advantages of fast imaging speed and low photon damage.

### Superior performance of DeepUI with noise and blur degradation

The physical degradation process brings superior performance of DeepUI. We choose SelfNet^27^ and 4BNet^18^, state-of-the-art isotropic methods using GAN architecture and 1D rotating super-resolution, respectively, for comparison. First of all, we conduct a simulation in different sample densities and lateral/axial similarity. We simulate the 3D point-shaped sample and the line-shaped sample with two densities. The three isotropic methods perform well in high similarity and sparse distribution situations, but SelfNet and 4BNet fail to realize isotropic results in dense distribution situations. In low-similarity line-shaped samples, SelfNet shows artifacts in both dense and sparse distributions. And 4BNet shows good performance in sparse distribution, but it fails to realize isotropic results in dense distribution. In contrast, DeepUI shows good performance in both situations. The quantitative metrics of RSP and RSE.

In real experiments, we choose the membrane of live vegetative *B. subtilis*, which shows discontinuity in the *xoz* plane after down-sampling, but it shows great axial and lateral similarity in Fig. 2(a). 4BNet shows line-shaped artifacts when conducting isotropic restoration in the absence of noise degradation. In contrast, SelfNet does not show obvious artifacts. But DeepUI, with a noise degradation step, shows better fidelity than SelfNet in the structure similarity index measure (SSIM) metrics in Fig. 2(b). The full width at half maximum (FWHM) of different methods is shown in Fig. 2(c). It shows that DeepUI and SelfNet have a better resolution than other methods, including the GT, 4BSIM. This is because 4BSIM can only realize near-isotropic performance with less axial frequency expansion compared with 4Pi-SIM.

**Figure 2.**
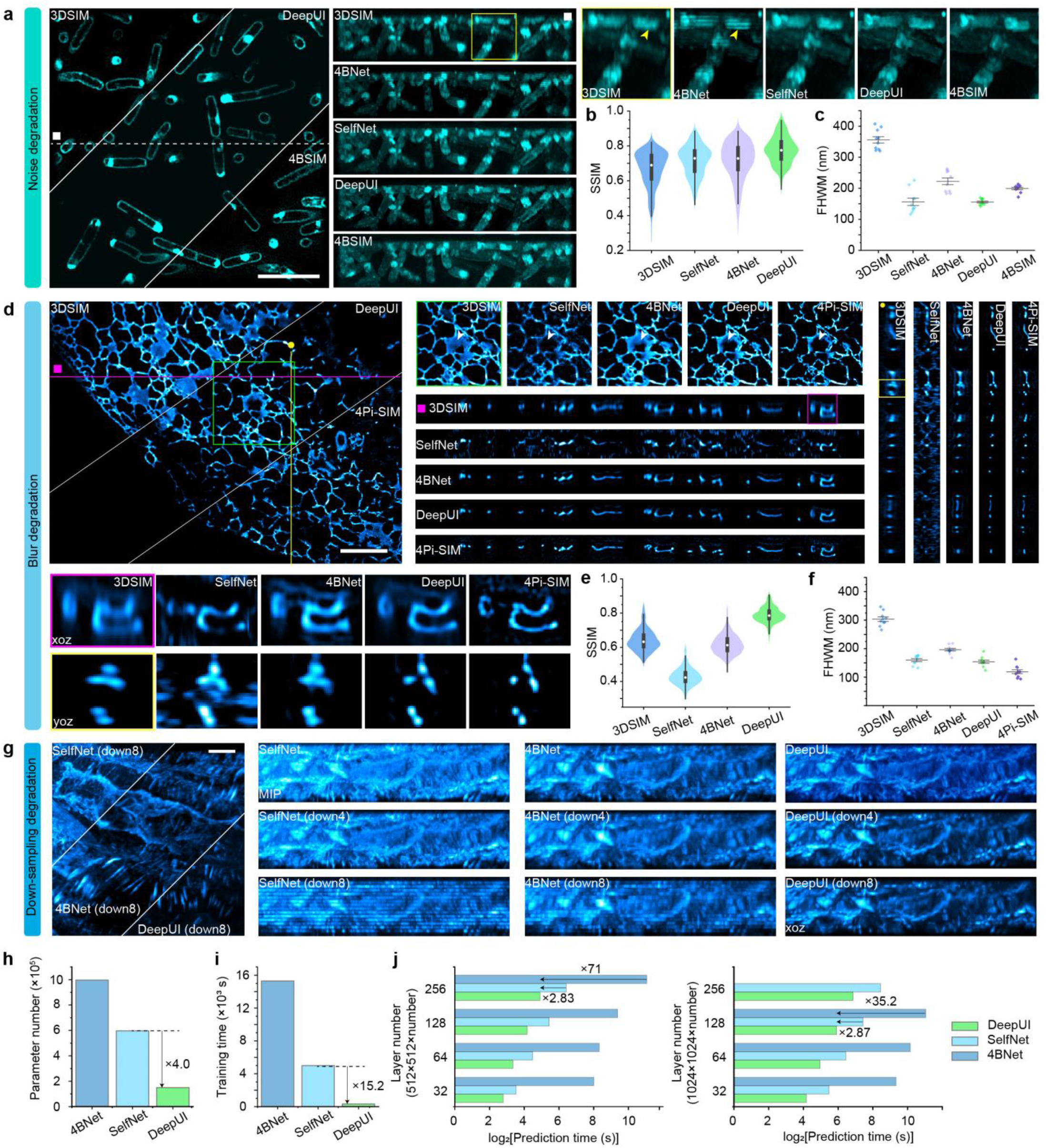
Comparison between DeepUI and other deep-learning-based isotropic methods to show the advantages of blur and noise degradation. (**a**) The advantages of noise degradation with the comparison of live vegetative *B. subtilis* between 3DSIM, DeepUI, 4BNet, SelfNet, and 4BSIM, with the corresponding (**b**) SSIM and (**c**) FWHM analysis. (**d**) The advantages of blur degradation with the comparison of ER between 3DSIM, DeepUI, 4BNet, SelfNet, and 4BSIM with the corresponding (**e**) SSIM and (**f**) FWHM analysis. (**g**) Comparison of DeepUI, SelfNet, and 4BNet in gradient axial down-sampling rate. The comparison of those methods when the axial scanning step is 8-fold than the lateral pixel size, namely 880 nm. The gradient axial scanning size from 110 nm to 880 nm, and the comparison of axial MIP images. The comparison of (**h**) parameter number, (**i**) training time, and (**j**) prediction time of different volume sizes, between DeepUI, 4BNet, and SelfNet. Scale bar: (**a, d**) 4 μm. (**g**) 8 μm.

Low similarity between the axial and lateral planes is also a characteristic of many other samples, for instance, subcellular thin samples. We chose the endoplasmic reticulum (ER) for illustration in Fig. 2(d). It shows DeepUI has a better correspondence of 4Pi-SIM in the lateral and axial planes. Its hollow structure in *xoy* images shows the most similarity to 4Pi-SIM. It can also distinguish the three point-shaped structures clearly, while other methods cannot. It’s noteworthy that SelfNet with a GAN network fails to learn the relationship because of the low similarity between the axial and lateral planes, resulting in many obvious artifacts. The SSIM and FWHM analysis in Fig. 2(e, f) also shows that DeepUI has better metrics.

The comparison of gradient axial down-sampling situations are shown in Fig. 3(g). Benefitting from the two-step network, DeepUI can greatly reduce the down-sampling artifacts and retrieve the 3D structure to a great extent, even in extreme conditions. In contrary, SelfNet and 4BNet fail to reconstruct the isotropic information in axial planes.

**Figure 3.**
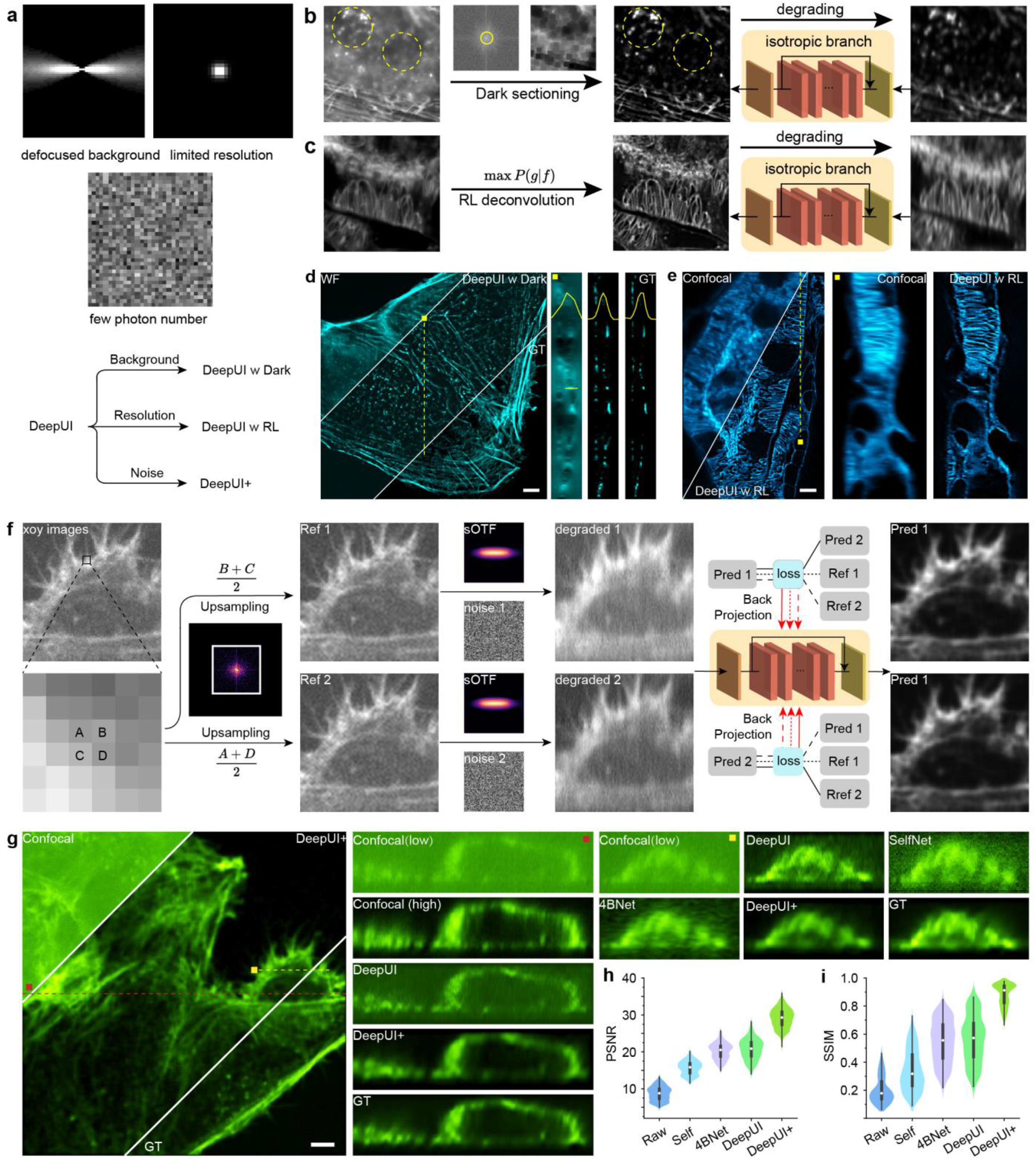
Extended DeepUI frameworks for joint de-backgrounding, deconvolution, and denoising tasks. (**a**) Three common barriers to decreasing the clarity of fluorescence images, including defocused background, limited resolution, and a photons number. DeepUI is developed as three extended methods to resolve those problems. (**b**) Integration with Dark sectioning in the pre-process step can improve the isotropic performance in the case of severely defocused background. (**c**) The comparison of the actin filament imaging results using DeepUI with or without Dark. The modified 3DSIM result without *xoy* super-resolution is used as ground-truth. (**d**) Integration with RL deconvolution in the pre-process step can improve the isotropic resolution. (**e**) The comparison of a mouse kidney section using DeepUI with or without RL. (**f**) The flowchart of the self-supervised denoised DeepUI, named DeepUI+. (**g**) Comparison of Confocal images with low light illumination, DeepUI, DeepUI+, SelfNet, 4BNet, and DeepUI+ with high light illumination of actin filament of Confocal imaging. The corresponding (**h**) PSNR and SSIM of those methods. Scale bar: 4 μm.

We further test the generality of DeepUI of different samples in a Confocal system. We use a subcellular (actin filament) model for testing two tissue samples (onion skin and mouse kidney), with the magnified zones. It can be seen that DeepUI can not only perform best in the cellular sample of the actin model, but also perform well in the tissue samples of onion skin and mouse kidney. Because of the non-similarity and low SNR, SelfNet exhibits artifacts, and 4BNet shows insufficient improvement in the axial resolution.

Because DeepUI simplified the flowchart from dual-GAN or 3D network to 2D RCAN network , DeepUI has greatly reduced network parameters (Fig. 2(g)), resulting in a greatly reduced training time (Fig. 2(h)) and prediction time (Fig. 2(i)). The parameter number and training time were reduced by about 4 and 15-fold, respectively. And the prediction time in various image sizes typically decreases about 3-fold compared with SelfNet, and 70-fold compared with 4BNet.

### Extension of DeepUI to different imaging tasks: de-background, deconvolution, and denoise

The results demonstrate the superior performance of DeepUI under challenging conditions such as low structural similarity, high information density, and axial down-sampling, surpassing existing state-of-the-art methods. However, several significant obstacles remain in practical applications^20,33,34^, most notably defocused background, inherent resolution limits, and low photon count scenarios. To address these limitations, we have developed three extended variants of our framework, as outlined in Fig. 3(a): DeepUI w Dark, DeepUI w RL, and DeepUI+.

Firstly, to eliminate the defocused background, we use Dark sectioning^33^ for pre-processing in Fig. 3(b). Dark sectioning, our previous work, is time and memory-efficient with high fidelity de-background performance. It can greatly remove the defocused background while maintaining weak information. Then, in the conditions where lateral resolution is not enough, we use Richardson-Lucy (RL) deconvolution^19^ for the pre-processing. RL deconvolution is a traditional and widely used method for improving the image resolution, and we here use it for further resolution improvement of DeepUI in Fig. 3(c). The actin filament imaged by WF microscopy before and after Dark sectioning is shown in Fig. 3(d). It can be shown that Dark sectioning can help DeepUI to obtain isotropic results with almost no backgrounds. And the modified 3DSIM result without the lateral resolution enhancement is used as GT, where DeepUI, with the help of Dark sectioning, shows great similarity with GT. The DeepUI results with RL deconvolution are shown in Fig. 3(e), indicating that RL can further improve the *xoy* and *xoz* resolution, with clearer visualization of the actin details. It is also noteworthy that DeepUI is different from the popular deconvolution methods. We chose the nuclear pore complex in and mouse kidney actin for the comparison of RL deconvolution and Sparse deconvolution. It shows that although deconvolution methods can achieve better 3D resolution, they fail to realize isotropic resolution with a 2∼4-fold anisotropic ratio. DeepUI can be directly applied to raw images to achieve isotropic resolution. Moreover, it can be effectively combined with deconvolution methods, where applying DeepUI to pre-deconvolved results further enhances their isotropy, as often 2D deconvolution cannot improve the anisotropy between lateral and axial resolution. It is important to note that preprocessing steps like Dark sectioning and RL deconvolution alter the effective PSF. Consequently, the *sOTF* must be recalibrated to ensure optimal performance.

Furthermore, we designed a denoising DeepUI method to improve the SNR following the zero-shot framework in Fig. 3(f). We use the spatial sampling of the nearest pixels to form four sub-images A, B, C, D^34-36^. Each sub-image is degraded into the simulated *xoz* images. Then, the degraded sub-images are randomly reconstructed by the DeepUI network, and the *xoy* images are used for the cross-validation. The sub-images are supervised by each other, and the isotropic performance is supervised by the *xoy* images. Then, the isotropic 3D stack can be obtained in Fig. 3(f), and we name it DeepUI+. To demonstrate the performance of DeepUI+, we use the high and low actin filament imaged by spinning-disk confocal microscopy for illustration in Fig. 3(g). Although the raw image has an extremely low SNR, DeepUI+ can greatly reduce the noise with high fidelity, as the GT shows, without compromise of the improved axial resolution. In this situation, SelfNet, 4BNet, and DeepUI will introduce the artifacts with noise, while DeepUI+ can maintain the good performance.

Due to the generality of DeepUI, we demonstrate that DeepUI+ can greatly reduce the artifacts in 3DSIM imaging caused by low SNR in actin filaments or motion artifacts of tubulin. We can also further explore the application of DeepUI in various state-of-the-art fluorescence super-resolution microscopy techniques, such as 3D-STED microscopy^16,37^ and 3DSIM.

## Discussion

We have introduced DeepUI, a physics-driven zero-shot framework that addresses the most persistent limitations in fluorescence microscopy: axial anisotropy and down-sampling. The core strength of our approach lies in its physically grounded and computationally efficient design. Unlike GAN-based or simulation-reliant methods, DeepUI explicitly models the anisotropic degradation process through a learned *sOTF* and realistic noise simulation. This ensures high fidelity and minimizes artifacts, particularly in challenging scenarios where axial and lateral information share low structural similarity, the simulated PSF does not correspond well to the actual situation, and unseen up-sampling noise, which often causes failure in other methods.

The advantages of DeepUI can be attributed to four key factors. First, the physics-guided blur degradation provides a stable and accurate prior for the restoration network, ensuring that the resolution enhancement adheres to physical constraints. Second, the explicit modeling of axial-specific noise enables robust performance under axial down-sampling and non-ideal imaging conditions, such as those with overwhelming noise or defocused backgrounds. Third, the framework’s design grants it remarkable generality, allowing it to handle complex samples, severe axial down-sampling, and diverse microscopy modalities—from high-end 3D-SIM and STED to conventional confocal systems. Finally, the framework’s extensibility, demonstrated by DeepUI w Dark, DeepUI w RL, and DeepUI+, allows it to be adapted to overcome common practical barriers like background fluorescence, limited native resolution, and low signal-to-noise ratio.

While DeepUI demonstrates high fidelity, the interpretability and reliability of deep learning outputs remain an important consideration in biological research. Future work could integrate uncertainty quantification methods, such as Bayesian deep learning, to provide confidence maps for the isotropic reconstructions. This would offer biologists valuable guidance on the reliability of fine structural details. Furthermore, for very thick specimens or large fields-of-view where the point spread function (PSF) varies significantly, a single *sOTF* might be insufficient. Adopting a block-wise processing strategy or developing a large model^28,38^ containing various *sOTF* could address this spatial variability. Lastly, the current framework employs a lightweight RCAN for efficiency. Replacing this with more advanced network architectures, such as DFCAN^39^, SRGAN^40^, diffusion models^41^ , or transformer-based networks, could potentially yield further improvements in reconstruction quality, albeit at a potential computational cost.

In conclusion, DeepUI achieves high-fidelity isotropic resolution from a single, anisotropic, and down-sampled 3D image stack without the need for paired training data or specialized hardware. Its exceptional speed—requiring only about 5 minutes for training and 0.5 minute for prediction—combined with its robustness and generality across various samples and state-of-the-art imaging techniques, makes it a highly practical solution. Given the significant technical complexity and cost associated with achieving isotropic resolution through optical approaches, DeepUI stands as a powerful yet accessible computational alternative, poised to advance multi-scale 3D imaging in the life sciences.

## Acknowledgments

This work was supported by the National Key R&D Program of China (2022YFC3401100), the National Natural Science Foundation of China (62025501, 62335008, 92150301, 62411540238, 624B2009, 62305004), and Major Basic Research Project of the Natural Science Foundation of Shandong Province (ZR2024ZD27). We thank the National Center for Protein Sciences and the Core Facilities of Life Sciences at Peking University. We would like to acknowledge the assistance of SLSTU-Nikon Biological Imaging Center at Tsinghua University. We thank the Optofem Technology Limited for providing MIRAVA Polyscope STED nanoscope (Abberior Instruments).

## Author Contributions Statement

R.C. and P.X. initiated and conceived the research. P.X. and M.L. supervised the project. R.C. developed the method and experiment. T. J. built the Fiji plugin and GUI. F. X. and K. Y. conducted the brain map and neuron experiment. Y. H., Y. L., W. R., and W. W. helped with the conception. Y. F., L. L., S. G., and D. S. conducted the live cell experiments. B. J. conducted the zebrafish experiment. G. X. conducted the live Drosophila experiment. H. W. helped with the figure drawing. R.C. drew the figures and videos. X. S. and D. K. provide experimental assistance. R. C., M.L., and P.X. wrote the manuscript with input from all authors.

## Methods

### Cell culture and label

For ER imaging, COS-7 cells were seeded onto μ-Slide eight-well chambers (ibidi, 80827). The ER was labeled by expression of Sec61β–mYonghong. After an 8 h incubation with the transfection mixture, the medium was replaced with standard growth medium before imaging.

### Open-source data and commercial sample

The tubulin, mitochondria, and membrane in Fig. 1(a, d), Fig. 2(a) are derived from 4B-SIM^18^. The ER in Fig. 2(d) is derived from 4Pi-SIM^13^. We also use some commercial standard samples for imaging^27^. The mouse kidney sections were purchased from Thermo Fisher (F24630). The onion skin was purchased from Suzhou Shenying Optical Co. The actin of COS-7 cells was purchased from GATTA (GATTA-Cells 4C).

### Imaging details

The Confocal imaging uses a Nikon AXR laser scanning microscope (60x objective, NA=1.4) to image actin filaments, onion skin, and mouse kidney. The spinning-disk Confocal imaging uses Ariy NovaSD microscope (60x objective, NA=1.4) to image neurons, mouse kidney, actin, and nematode. The 3DSIM imaging uses GE | OMX microscope (60x objective, NA=1.4) to image ER, synaptonemal complexes, and tubulin.

### Training process

The pre-training of degradation uses a UNet network^43^ to exclude the influence of initial *sOTF*. The Unet contains three encoder and decoder blocks with 15 convolution layers and 3 concat blocks, following the traditional Unet framework. The activation function is ReLu, and the learning rate is 1e-4 for all experiments. Typically, one epoch can obtain the ideal results. For one system with one certain wavelength, the *sOTF* does not need to be calculated for each sample with a similar thickness.

The training of DeepUI and DeepUI follows the RCAN network. The RCAN network implemented for this study utilizes a streamlined architecture with 2 residual groups, each containing 3 residual channel attention block, for deep feature extraction. The activation function is ReLu, and the learning rate is 1e-3 for all experiments. Typically, one to three epochs can obtain the ideal results.

The dataset generation step uses MATLAB 2021b, and training uses PyCharm (Pytorch 2.6.0, CUDA 12.6). The framework is light with no strict requirements for a computer. We train and predict the model with a common computer equipped with an Intel i7-12700 CPU and a NVIDIA GeForce GTX 1650.

### Calibration of RSP, RSE, and FWHM

We use the PSNR and SSIM to evaluate the similarity between the calculated image and the GT image. The PSNR and SSIM are written by the function in MATLAB.NanoJ-SQUIRREL^44^ is used to quantify the difference between the reconstructed images when GT is available. RSE means the sum of error between the scaled GT image and reconstructed image, and RSP represents the Pearson coefficient between the scaled GT image and reconstructed image. The closer the RSP is to 1, the closer the RSE is to 0, indicating that the GT and reconstructed images are more similar. In order to quantify the axial resolution, we use FWHM. The profiles of the images along the *z*-axis are plotted, and Gaussian fitting is conducted. The FWHM of Gaussian curves is calculated at 5 or 0 different places.

### Imaging process and analysis

We use Fiji to conduct MIP in the *xoy*/*xoz* plane, FWHM calculation, and RSP/RSE calculation^44^. We use MATLAB (2021b) to conduct the SSIM, PSNR^45^. The segmentation and surface analysis use the Imaris (10.1.0). Origin (OriginLab, V2021) is used to plot curves, and the brightness and contrast are adjusted linearly for display purposes.

